# Metabolomics-Guided Machine Learning Reveals Diagnostic and Mechanistic Biomarkers in CHB with MASLD

**DOI:** 10.1101/2025.08.19.671181

**Authors:** Chuyang Wang, Yutao Chen, Huanming Xiao, Jianxiong Cai, Ruihua Wang, Xuan Zeng, Ming Lin, Wofeng Liu, Xiaoling Chi, Qubo Chen

## Abstract

**Background:** Metabolic dysfunction–associated steatotic liver disease (MASLD) often coexists with chronic hepatitis B (CHB), yet early diagnosis remains challenging, particularly in non-obese patients or those with subclinical features. This study aimed to identify metabolic signatures of CHB-related MASLD and construct a predictive model using untargeted metabolomics integrated with machine learning.

**Methods:** Serum metabolomics was performed on 160 subjects (80 CHB + MASLD and 80 healthy controls). Differential metabolites were identified and analyzed using KEGG enrichment and 4 machine learning algorithms (Random Forest, XGBoost, SVM, and LASSO). Metabolite–clinical correlations and diagnostic model performance were evaluated.

**Results:** A total of 924 differential metabolites were identified, with significant enrichment in pathways related to the TCA cycle, sphingolipid metabolism, and amino acid turnover. Machine learning prioritized six robust and biologically relevant metabolites: L-aspartic acid, succinic acid, caproic acid, sebacic acid, monomenthyl succinate, and glycolaldehyde, which consistently distinguished CHB + MASLD patients from controls (AUC > 0.75). These metabolites reflect key disruptions in mitochondrial function, lipid oxidation, and redox homeostasis. Integrated models combining metabolomics with clinical indices achieved perfect classification (AUC = 1.000).

**Conclusion:** CHB-associated MASLD is driven by systemic metabolic remodeling centered on mitochondrial overload, oxidative stress, and impaired amino acid metabolism. The identified metabolites provide mechanistic insights and hold promise for non-invasive MASLD screening in CHB patients. This study underscores the potential of multi-algorithmic metabolomics in advancing early diagnosis and personalized management of complex liver comorbidities.

## Introduction

Chronic hepatitis B (CHB), caused by persistent infection with the hepatitis B virus (HBV), remains a major global health burden. According to the World Health Organization, approximately 296 million individuals were living with chronic HBV infection in 2019, with around 820,000 deaths attributable to HBV-related liver failure, cirrhosis, and hepatocellular carcinoma (HCC) that year^1^. China bears a disproportionate share of this burden, with an estimated hepatitis B surface antigen (HBsAg) prevalence of 6.1% in 2016, accounting for nearly one-quarter of global cases^2^. Although long-term antiviral therapy with nucleos(t)ide analogs such as entecavir or tenofovir has been shown to reduce the risk of HCC in CHB patients, it does not fully eliminate the threat^3^.

In parallel, metabolic dysfunction–associated steatotic liver disease (MASLD), formerly known as non-alcoholic fatty liver disease (NAFLD), has emerged as another leading cause of chronic liver injury. Recognizing that liver steatosis often occurs in the context of obesity, type 2 diabetes, and other cardiometabolic disorders, international liver societies recently endorsed the term “MASLD” to better reflect its metabolic roots^4–6^. Given the increasing rates of obesity, hypertension, hyperlipidemia, and insulin resistance worldwide^7,8^, the prevalence of MASLD is expected to continue rising.

The intersection of CHB and MASLD represents a complex and increasingly common clinical scenario. Previous studies on CHB metabolomics have mostly focused on patients without hepatic steatosis, while patients with CHB combined with MASLD have a poorer prognosis^9^. Early studies have shown that patients with CHB combined with NAFLD have an increased risk of HCC^10,11^. Risk factors of MASLD, such as cardiovascular metabolic risk factors such as diabetes, may increase the risk of HCC^12^. There are few reports on the impact of the coexistence of MASLD and CHB on the risk of HCC in CHB patients. However, the biological mechanisms underlying the comorbidity of chronic hepatitis B (CHB) and metabolic dysfunction-associated steatotic liver disease (MASLD) remain poorly characterized. In particular, the molecular crosstalk between viral hepatitis-induced liver injury and metabolic dysregulation, is not well understood, and only limited studies have addressed their integrated metabolic profiles. Therefore, analyzing the changes in serum metabolites of CHB combined with MASLD patients through modern metabolomics techniques not only helps to elucidate the modern biological mechanisms of the disease, but also provides a basis for comprehensive treatment.

Metabolomics offers a powerful approach to explore disease mechanisms through the comprehensive profiling of small-molecule metabolites. By capturing systemic metabolic alterations, metabolomics provides insight into both host physiology and pathophysiological processes at the molecular level^13,14^. Liquid chromatography– tandem mass spectrometry (LC-MS/MS) has become one of the most widely used technologies in this field, offering high sensitivity, broad coverage, and compatibility with diverse metabolite classes, especially polar and thermally unstable compounds^15–17^.

In addition, studies have linked lower levels of branched-chain amino acids (BCAAs) with MASLD severity and insulin resistance^18^. However, the metabolic signature of CHB patients co-diagnosed with MASLD has not yet been well defined, and understanding this profile could be crucial for early diagnosis, risk stratification, and therapeutic targeting.

Therefore, we hypothesize that CHB-MASLD patients exhibit a unique metabolic signature that reflects the synergistic impact of viral infection and metabolic dysfunction. Identifying these metabolomic alterations may improve our understanding of disease mechanisms and reveal novel diagnostic or therapeutic targets.

In this study, we performed untargeted metabolomic profiling of serum samples from patients with CHB and coexisting MASLD, as well as healthy controls. Using LC-MS/MS combined with multivariate statistical analysis, we identified significantly altered metabolites and enriched metabolic pathways. We further applied machine learning algorithms, including Random Forest, XGBoost and SHAP, SVM with SHAP, and LASSO logistic regression to identify key metabolites with high discriminative power. In addition, we assessed the relationships between selected metabolites and clinical parameters to uncover potential biological relevance. Our findings provide new insights into the metabolic disturbances underlying CHB-MASLD and offer candidate biomarkers for improved diagnosis and therapeutic decision-making.

## Material and method

### Sample collection and grouping

This study enrolled a total of 160 participants, including 80 patients diagnosed with chronic hepatitis B (CHB) coexisting with metabolic dysfunction-associated steatotic liver disease (MASLD), and 80 age- and sex-matched healthy controls. The diagnosis of CHB was based on the persistence of hepatitis B surface antigen (HBsAg) positivity for more than six months, along with detectable HBV DNA and/or abnormal liver function tests. MASLD was diagnosed by imaging-confirmed hepatic steatosis (via ultrasound, MRI, or CT) combined with at least one cardiometabolic risk factor, including BMI ≥23 kg/m², elevated fasting blood glucose (≥5.6 mmol/L) or known type 2 diabetes, hypertension (≥130/85 mmHg), hypertriglyceridemia (TG ≥1.70 mmol/L), or low HDL-C (≤1.0 mmol/L for males, ≤1.3 mmol/L for females). Exclusion criteria included other viral hepatitis (e.g., HCV, HDV), significant alcohol consumption (>210 g/week for men or >140 g/week for women), autoimmune liver diseases, and incomplete clinical records. The diagnosis of CHB+MASLD was based on established clinical guidelines. Fasting venous blood samples were collected from each participant, and serum was separated by centrifugation at 3,000 rpm for 10 minutes and stored at – 80°C until further analysis.

Serum samples had been collected previously from patients with CHB+MASLD at the Department of Hepatology, Guangdong Provincial Hospital of Traditional Chinese Medicine between November 2021 and October 2022, and from healthy controls at the Physical Examination Center of Guangzhou Eleventh People’s Hospital between May 2022 and August 2023. For the present study, archived serum samples were accessed in September 2023 for retrospective analysis. All samples were anonymized before analysis, and investigators did not have access to personally identifiable information during or after data collection.

All participants had provided written informed consent at the time of sample collection. The research protocol for the retrospective use of these archived samples was reviewed and approved by the Ethics Committee of Guangdong Provincial Hospital of Traditional Chinese Medicine (Approval No.: ZE2023-194-01).

### Metabolome data sequencing and preprocessing

Perform metabolite detection on all serum samples using LC-MS/MS. Serum samples were processed using protein precipitation and extracted with methanol, followed by centrifugation and filtration. Metabolite detection was carried out in both positive and negative electrospray ionization (ESI) modes. Raw data were preprocessed, including peak detection, alignment, and annotation. Metabolomic data were processed using Progenesis QI software. After feature alignment and metabolite annotation using the Metlin™ MS/MS library, the following preprocessing steps were applied: (1) outlier filtering based on the coefficient of variation (CV); (2) feature filtering by retaining only those with <50% missing values in any group; (3) missing value imputation using half of the minimum detected value; and (4) normalization based on the total ion current (TIC) per sample. This approach minimized technical variability and allowed for reliable comparison across samples. Data from positive and negative ion modes were then merged for integrated analysis. Perform partial least squares discriminant analysis (PLS-DA) on metabolic profile data through R package (ropls, v_1.34.0) for preprocessing and pattern recognition.

### Differential Metabolite Analysis

To determine the metabolites that showed significant changes between the CHB+MASLD group and the control group, a combination of univariate and multivariate statistical analyses was used. The screening criteria for differential metabolites are: VIP>1, | log ₂ FC |>0.01, and p<0.05. Perform metabolite differential analysis using the MetaboAnalyst platform (https://www.metaboanalyst.ca/home.xhtml) and perform KEGG pathway enrichment analysis on differential metabolites.

### XGBoost with SHAP Interpretation

To evaluate the diagnostic utility of selected metabolites, we constructed an Extreme Gradient Boosting (XGBoost) classification model using the top 22 metabolites identified from the random forest analysis. The model was trained on all samples using five-fold cross-validation. Model performance was evaluated based on the area under the receiver operating characteristic curve (AUC). SHapley Additive exPlanations (SHAP) values were used to interpret the contribution of individual metabolites to the prediction outcome. SHAP summary and dependence plots were generated to highlight the most impactful features. The above analysis was conducted in R, including the R package: xgboost,v_1.7.11.1, randomForest(v_4.7-1.2), pROC(v_1.18.5),SHAPforxgboost(v_0.1.3).

### SVM with SHAP Interpretation

Support Vector Machine (SVM) models were also constructed using the same set of 22 metabolites to validate findings from XGBoost(R package:e1071, v_1.7-16). The radial basis function (RBF) kernel was applied, and five-fold cross-validation was used to assess model generalizability. SHAP analysis was employed to interpret metabolite contributions to classification. Top-ranking metabolites identified by SHAP were further compared between groups, and their relative abundance was visualized.

### LASSO Logistic Regression

To further refine biomarker selection, we performed least absolute shrinkage and selection operator (LASSO) logistic regression using the glmnet package in R(v_4.1-8). The model was trained on a matrix of 22 metabolite expression values, with CHB+MASLD versus control as the binary outcome variable. Ten-fold cross-validation was conducted to determine the optimal regularization parameter (lambda). Metabolites with non-zero coefficients at lambda.min were selected as key discriminative features. Model performance was assessed by ROC analysis.

### Correlation analysis of clinical parameters and construction of a combined logistic regression model

To explore the potential clinical relevance of the selected metabolites, Spearman correlation analysis was conducted between the 22 key metabolites and 7 clinical parameters: body mass index (BMI), alanine aminotransferase (ALT), aspartate aminotransferase (AST), triglycerides (TG), total cholesterol (TC), low-density lipoprotein cholesterol (LDL-C), and high-density lipoprotein cholesterol (HDL-C). Correlation matrices were visualized as heatmaps. Additionally, a combined logistic regression model integrating both clinical parameters and metabolite features was constructed and evaluated using ROC and cross-validation.

## Results

### Global Metabolomic Profiling Reveals 924 Differential Metabolites in CHB + MASLD

Untargeted metabolomics was performed on 160 serum samples (80 CHB + MASLD, 80 healthy controls, Figure 1A). PLS-DA indicated a clear separation of the two groups, underscoring a distinct metabolic signature (Figure 1B). Using VIP > 1, |log₂FC| > 0.01 and *p* < 0.05, 924 metabolites were significantly altered, of which 311 were up-regulated and 613 down-regulated in CHB + MASLD (Figure 1C). KEGG enrichment highlighted sphingolipid metabolism, pyruvate metabolism, nicotinate and nicotinamide metabolism, histidine metabolism, glyoxylate and dicarboxylate metabolism, glycine, serine and threonine metabolism, TCA cycle, arginine biosynthesis alanine, aspartate and glutamate metabolism (Figure 1D). Random-forest ranking (MeanDecreaseAccuracy) further reduced the panel to 22 biologically meaningful metabolites, monomenthyl succinate, sphinganine, arachidonic acid, succinic acid, L-aspartic acid and sebacic acid,supported by concordant MeanDecreaseGini scores (Figure 1E–F). These 22 candidates were passed to XGBoost, SVM and LASSO logistic for in-depth feature interpretation.

**Figure 1.**
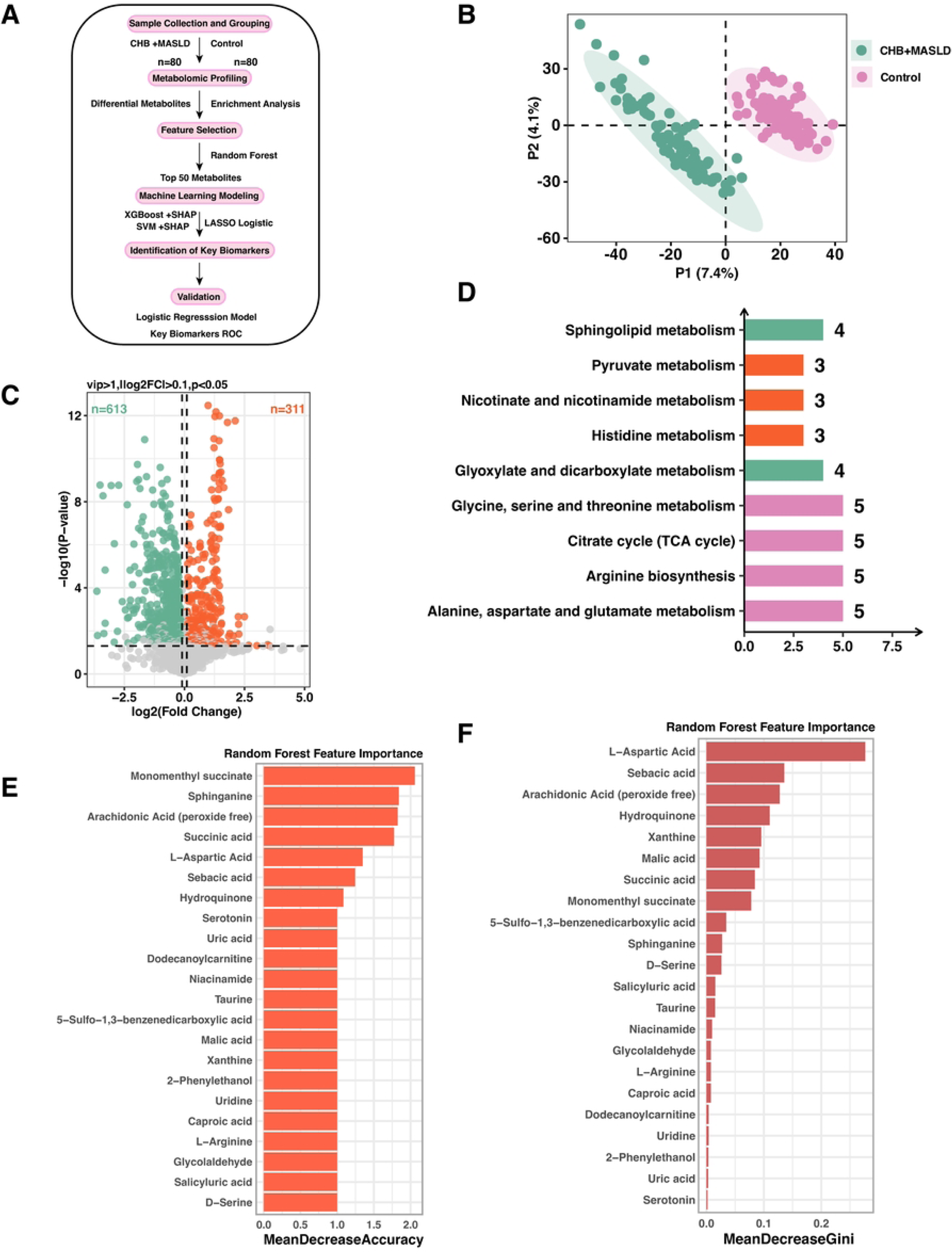
Overview of metabolomics analysis and identification of key metabolites. **A.** Schematic workflow of the study. A total of 160 serum samples were collected, including 80 from CHB combined with MASLD patients and 80 from healthy controls. Untargeted metabolomics was performed, followed by differential metabolite analysis and KEGG enrichment. The top 50 metabolites were selected via Random Forest based on MeanDecreaseAccuracy, and then XGBoost+SHAP and SVM+SHAP models were constructed to identify key metabolites. These were subsequently combined with clinical indicators for logistic regression modeling, and final candidate metabolites were validated by ROC analysis. **B.** PLS-DA plot demonstrating clear separation between the CHB+MASLD group and the control group. **C.** Volcano plot of differential metabolites based on VIP > 1, |log2FC| > 0.01, and p < 0.05. **D.** KEGG enrichment of differential metabolites showing significantly enriched pathways (p < 0.05). **E.** Metabolites selected by Random Forest based on MeanDecreaseAccuracy. Deleted metabolites without HMDB numbers and related to drug metabolism. **F.** Feature importance of the metabolites based on Random Forest MeanDecreaseGini values.

### XGBoost Identifies High-Impact Metabolites: L-Aspartic Acid, Monomenthyl succinate and sebacic acid

XGBoost modelling of the 22 metabolites yielded excellent discrimination (Figure 2A, AUC = 1.000); five-fold cross-validation remained robust (mean AUC = 0.9953; Figure 2B). Feature importance and SHAP plots (Figure 2C–E) pinpointed L-aspartic acid, monomenthyl succinate, sebacic acid, caproic acid and glycolaldehyde as the most influential variables. SHAP values indicated sebacic acid drove predictions toward CHB + MASLD, whereas L-aspartic acid and glycolaldehyde favoured the control class. Box plot comparisons confirmed higher levels of L-aspartic acid, monomenthyl succinate, caproic acid, arachidonic acid and glycolaldehyde in controls, but elevated sebacic acid in CHB + MASLD (*p* < 0.001; Figure 2F), supporting their discriminatory potential. Although the model showed near-perfect classification performance, the use of five-fold cross-validation mitigated the risk of overfitting and supports the robustness of the findings.

**Figure 2.**
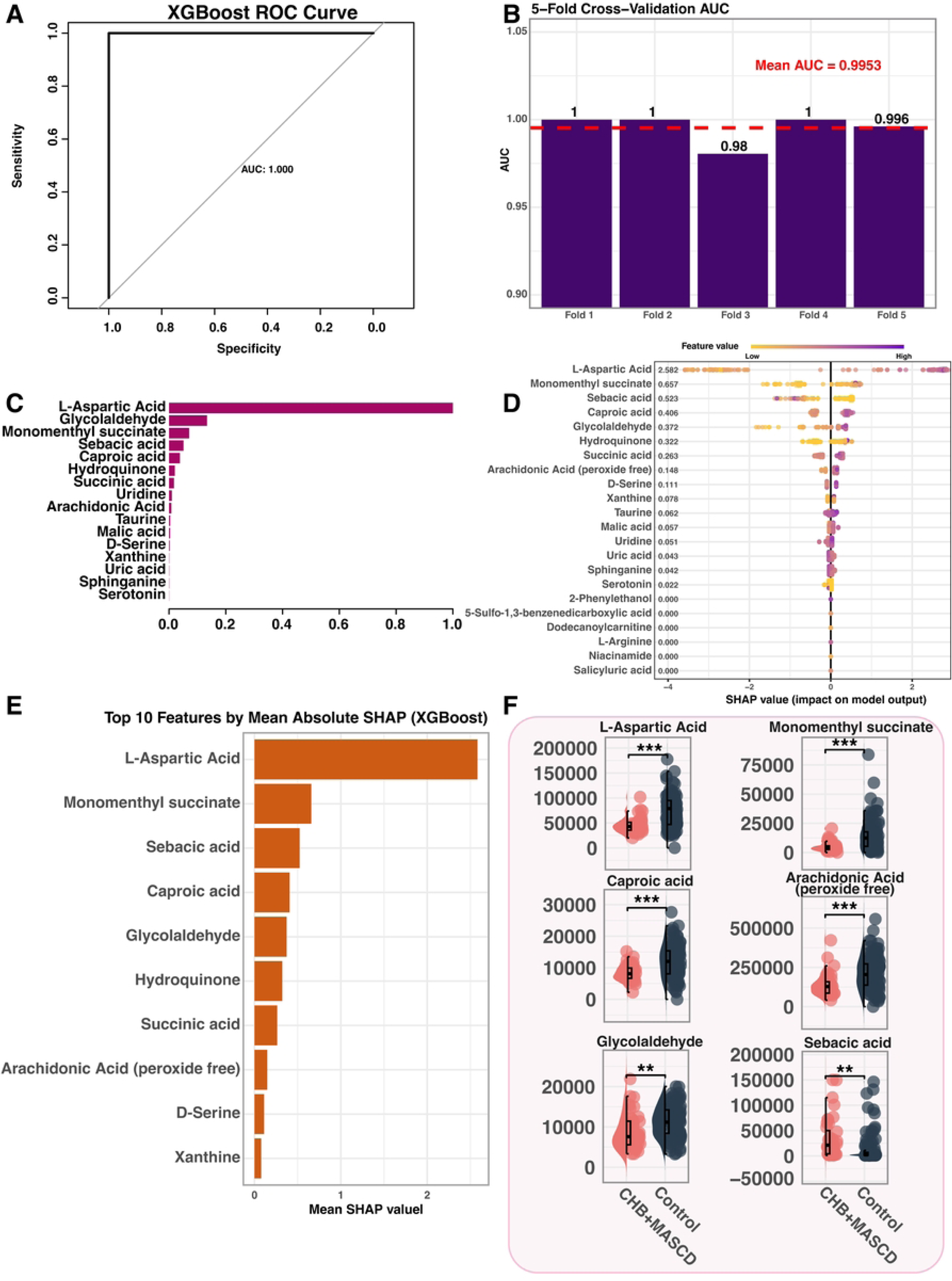
XGBoost-based feature selection and interpretation of differential metabolites. **A.** ROC curve of the XGBoost model. **B.** 5-fold cross-validation AUC of the model. **C.** XGBoost feature importance ranked by gain.. **D.** SHAP summary plot of the top 15 features. SHAP values: CHB+MASLD group (left), Control group (right). **E.** Bar plot of top 10 metabolites by mean absolute SHAP value in the XGBoost model. **F.** Comparison of relative abundance for 4 selected metabolites between CHB+MASLD and Control groups(**p<0.01,***p < 0.001).

### SVM Yields Concordant Classification and Highlights Additional Markers

The SVM model performed equally well (Figure 3A, AUC = 1.000; mean CV-AUC = 0.9984, Figure 3B) and achieved perfect sensitivity and specificity (Figure 3C). SHAP analysis revealed 20 metabolites such as L-aspartic acid, succinic acid, caproic acid, D-serine, sphinganine, uric acid and glycolaldehyde as top contributors (Figure 3D). Quantitative comparison showed 11 metabolites were significantly elevated in CHB + MASLD, whereas three were reduced (*p* < 0.001; Figure 3E), indicating potential pathophysiological relevance.

**Figure 3.**
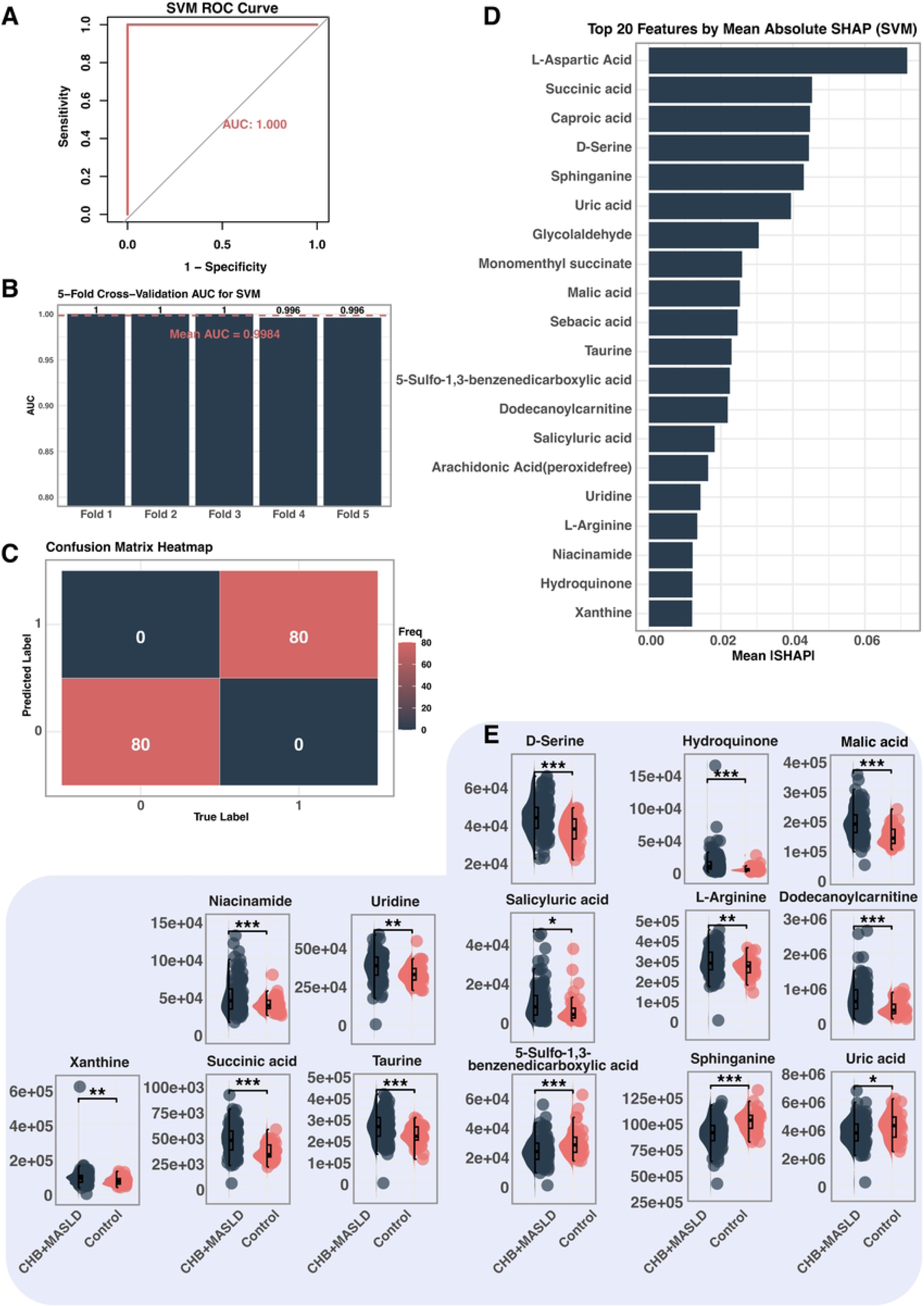
SVM-based feature selection and interpretation of differential metabolites. **A.** ROC curve of the SVM model. **B.** 5-fold cross-validation AUC values. **C.** Confusion matrix heatmap of predicted vs true labels. The model achieved perfect classification: 80 true positives (class 0), 80 true negatives (class 1), with no misclassifications. **D.** Top 20 features ranked by mean absolute SHAP value from the SVM model. **E.** Relative abundance of selected key metabolites. All displayed metabolites were significantly higher in the CHB+MASLD group compared to the Control group (*p<0.05, **p<0.01,***p < 0.001).

### LASSO Logistic Regression Narrows the Panel to Seven Core Metabolites

LASSO regression progressively shrank coefficients as the regularisation term increased (Figure 4A). Ten-fold cross-validation selected the optimal λ corresponding to the minimum mean cross-validated error (λ.min; Figure 4B), resulting in a model that retained seven non-zero metabolites: caproic acid, glycolaldehyde, monomenthyl succinate, D-serine, L-aspartic acid, succinic acid, and salicyluric acid (Figure 4C). We chose λ.min rather than the more conservative λ.1se in order to retain a broader panel of candidate metabolites with potential biological relevance for downstream validation and mechanistic exploration. However, we acknowledge that this choice may reduce model sparsity and compromise generalizability. The resulting model achieved excellent classification performance, with an AUC of 1.000 (Figure 4D), highlighting the diagnostic potential of this metabolite subset.

**Figure 4.**
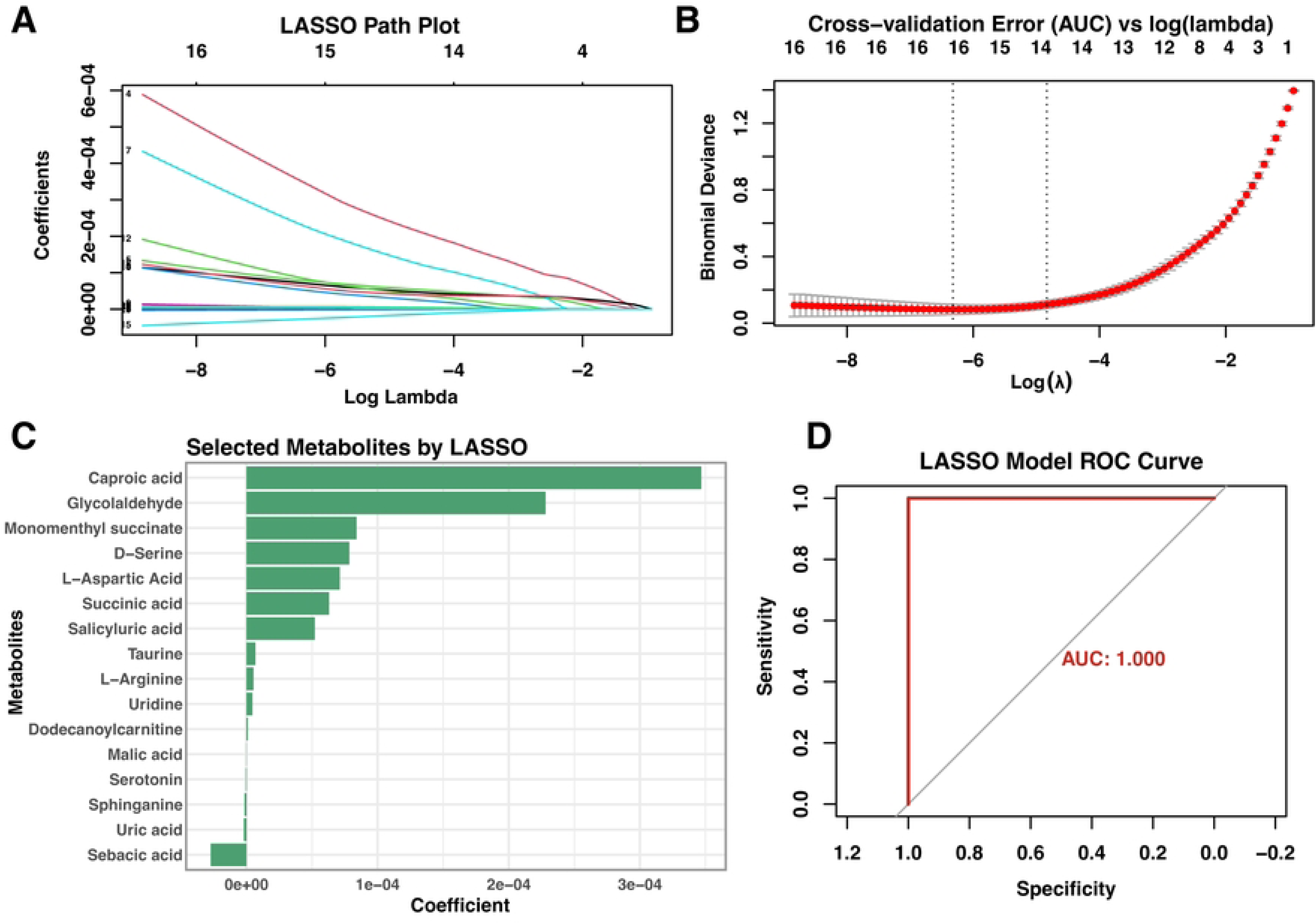
LASSO logistic-Based Feature Selection and Model Performance Evaluation. **A.** Path coefficient trajectory plot of LASSO logistic regression model, showing how coefficients of metabolites change with varying penalization strength. **B.** Cross-validation error plot for LASSO model tuning, with the optimal λ value minimizing classification error. **C.** Barplot of selected metabolites based on non-zero LASSO coefficients, representing their relative contribution to the model. **D.** ROC curve evaluating the diagnostic performance of the LASSO model, demonstrating high discriminative ability.

### Metabolite–Clinical Interplay and Integrated Modelling

Clinical comparison revealed significantly altered metabolic indicators, including increased BMI, ALT, AST, LDL-C, TC and TG, and decreased HDL-C in the CHB + MASLD group. Statistical significance was determined by t-tests and adjusted using FDR correction to control for multiple comparisons (Figure 5A). Notably, L-aspartic acid showed a significant positive correlation with HDL-C, while being negatively correlated with BMI, ALT, AST, LDL-C, TC, and TG. As a key amino acid involved in the urea cycle and transamination^19^, lower L-aspartic acid levels in metabolically dysregulated patients may reflect compromised hepatic amino acid metabolism and detoxification capacity, both of which are hallmarks of liver dysfunction. Caproic acid, a medium-chain fatty acid^20^, exhibited a similar pattern, which may suggest impaired β-oxidation and mitochondrial stress, commonly observed in fatty liver conditions.

**Figure 5.**
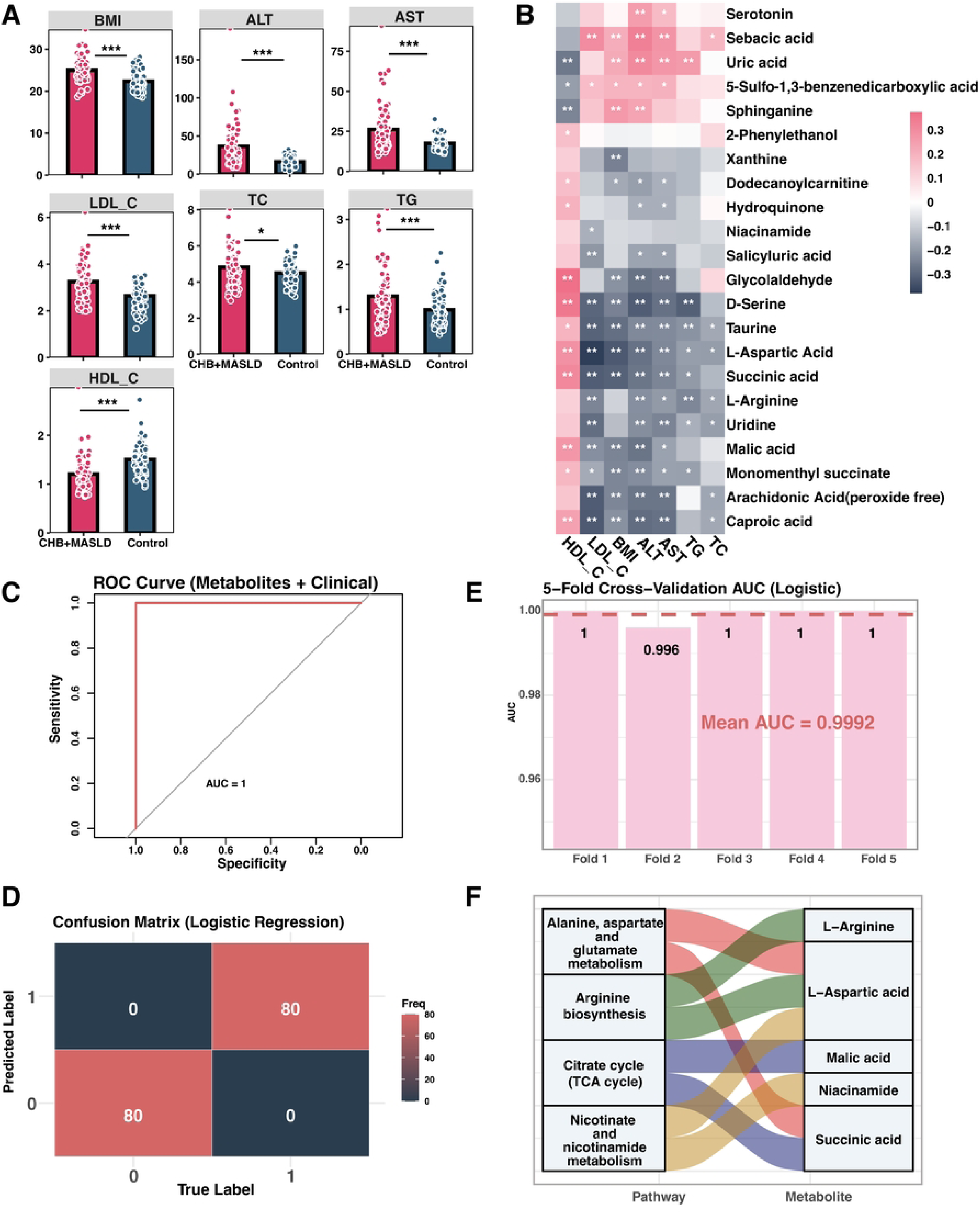
Clinical characteristics, metabolite–clinical associations, and logistic regression modeling. **A.** Comparison of clinical parameters between CHB+MASLD and control groups(*p < 0.05, **p < 0.01, ***p < 0.001). **B.** Heatmap of Spearman correlations between 15 key metabolites and 7 clinical indicators (*p < 0.05, **p < 0.01, ***p < 0.001). Pink indicates positive correlation, while dark gray indicates negative correlation. **C.** ROC curve for the logistic regression model constructed using the 15 metabolites plus clinical indicators. **D.** Confusion matrix for the logistic regression model. **E.** 5-fold cross-validation AUCs of the logistic regression model. **F.** KEGG Pathway Enrichment of 22 Key Metabolites with Metabolite-Pathway Mapping. KEGG enrichment analysis of the 22 key metabolites identified through multiple machine learning models. Only significantly enriched pathways (p < 0.05) are shown. A chord diagram illustrates the association between each enriched pathway (left) and the metabolites involved (right), highlighting overlapping metabolites across different biological pathways.

Glycolaldehyde, a reactive aldehyde involved in oxidative glucose metabolism^21^, was also positively associated with HDL-C and negatively correlated with BMI, ALT, and AST. Its reduction may reflect increased oxidative stress and impaired aldehyde detoxification in CHB + MASLD. Additionally, monomenthyl succinate and D-serine were significantly positively correlated with HDL-C and negatively associated with all other clinical indicators, except for TC. D-serine is a co-agonist of the NMDA receptor and may reflect neurometabolic regulation and glutathione pathway balance^22^, while monomenthyl succinate an ester derivative of succinic acid may indicate TCA cycle alterations^23^, often dysregulated in metabolic liver disease. Collectively, these correlations suggest that specific metabolites are tightly linked to hepatic function, lipid metabolism, and inflammatory burden. The inverse relationships with BMI and liver enzymes support their potential role as markers of metabolic homeostasis or disruption in CHB with MASLD (Figure 5B). A combined logistic model incorporating all 22 metabolites and seven clinical variables achieved an AUC of 1.000, perfect accuracy in the confusion matrix, and a mean CV-AUC of 0.996 (Figure 5C–E), confirming the additive diagnostic value of metabolic information. While these results demonstrate excellent discriminative performance, we acknowledge the possibility of overfitting due to the high dimensionality of inputs relative to sample size. Nevertheless, five-fold cross-validation was used throughout to mitigate this risk, and future validation in independent cohorts will be essential to confirm generalizability.

KEGG enrichment of the 22 metabolites yielded four significant pathways (p < 0.05): Alanine, aspartate and glutamate metabolism (L-aspartic acid, succinic acid), Arginine biosynthesis (L-arginine, L-aspartic acid), Citrate cycle (TCA cycle) (malic acid, succinic acid), and Nicotinate and nicotinamide metabolism (L-aspartic acid, niacinamide) (Figure 5F). These pathways converge on five metabolites—L-arginine, L-aspartic acid, malic acid, niacinamide, and succinic acid—linking amino acid turnover, energy production, and NAD⁺ metabolism, thereby highlighting their central role in CHB + MASLD pathophysiology.

### Single-Metabolite ROC Analysis Supports Diagnostic Potential

Individual ROC analysis of the 22 metabolites showed outstanding performance for L-aspartic acid, caproic acid, sebacic acid, arachidonic acid and hydroquinone (AUC > 0.90), strong performance for monomenthyl succinate, malic acid, D-serine, dodecanoylcarnitine and 5-sulfo-1,3-benzenedicarboxylic acid (AUC > 0.80), and acceptable accuracy for the remaining metabolites (AUC > 0.60) (Figure 6). These analyses were performed between the CHB + MASLD group and healthy controls. Grouping was based on comprehensive clinical evaluation, imaging findings, laboratory tests, and virological screening. These results further substantiate the diagnostic utility of the selected metabolites in distinguishing CHB patients with coexisting MASLD from healthy individuals.

**Figure 6.**
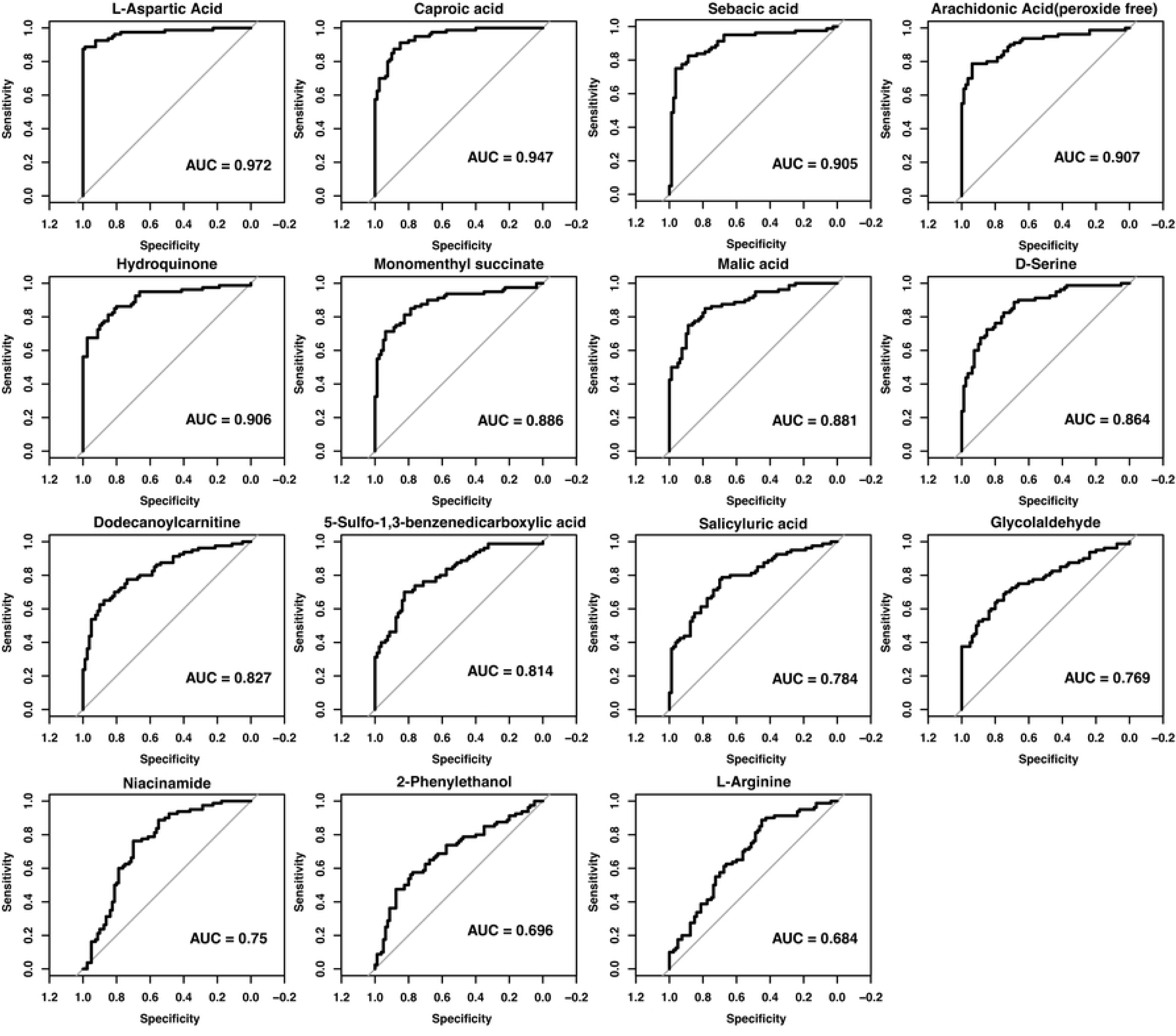
ROC validation of the 22 key metabolites.

## Discussion

### Metabolic Pathway-Level Disruption Reveals Lipid, Amino Acid, and Energy Metabolism as Central Axes in CHB + MASLD

Rather than isolated metabolite shifts, CHB + MASLD presents as a systemic metabolic reprogramming state, involving lipid overload, impaired nitrogen disposal, and mitochondrial dysfunction. KEGG enrichment of both the global and feature-selected metabolite sets consistently pointed to alterations in the tricarboxylic acid (TCA) cycle, amino acid metabolism, and lipid oxidation pathways, underscoring these axes as central to disease pathophysiology^24^. Notably, succinic acid and malic acid, two key TCA intermediates, were significantly elevated in CHB + MASLD patients^25,26^. This pattern suggests a potential metabolic bottleneck in oxidative phosphorylation, where substrate accumulation reflects either enhanced anaplerosis (e.g., from amino acids or fatty acids) or impaired downstream utilization by the electron transport chain^27,28^. Accumulation of TCA intermediates such as succinate has been linked to pseudohypoxic signaling through stabilization of HIF-1α, promoting hepatic inflammation and fibrosis progression^27^.

In contrast, L-aspartic acid, which links the TCA cycle to urea cycle function and amino acid nitrogen disposal, was markedly decreased. This depletion may indicate compromised mitochondrial aspartate export or impaired aminotransferase activity, leading to inefficient ammonia detoxification and contributing to nitrogen stress in the steatotic and virally inflamed liver^19,29^. Furthermore, enrichment of lipid related pathways, notably sphingolipid metabolism and markers of peroxisomal ω-oxidation such as sebacic acid points to lipid overload^30,31^.

Taken together, these findings depict CHB + MASLD as a syndrome of mitochondrial overload, impaired redox and nitrogen homeostasis, and adaptive but ultimately maladaptive lipid oxidation responses.

### Core Discriminatory Metabolites Implicate Mitochondrial Dysfunction, Lipid Stress, and Redox Imbalance in CHB + MASLD

Through an integrative approach combining three machine learning algorithms: XGBoost, SVM, and LASSO logistic regression. We identified six metabolites that consistently ranked among the top contributors to classification accuracy between CHB + MASLD patients and controls(STable 1). L-aspartic acid, caproic acid, and glycolaldehyde were prioritized across all three models, while monomenthyl succinate, D-serine, and succinic acid were identified by at least two models. These molecules are not only statistically robust but also biologically meaningful, as they cluster within key metabolic pathways involving mitochondrial function, lipid oxidation, and redox homeostasis.

L-aspartic acid is a multifunctional amino acid involved in the tricarboxylic acid (TCA) cycle, urea cycle, and nucleotide biosynthesis^19,32^. Its significant depletion in CHB + MASLD patients may reflect impaired mitochondrial export or dysfunction in the glutamate–aspartate shuttle^19^. Correlation analysis further supports its clinical relevance: L-aspartic acid levels were positively correlated with HDL-C and negatively correlated with ALT, AST, LDL-C, and TG, indicating a protective metabolic profile inversely associated with liver injury and dyslipidemia. Caproic acid (hexanoic acid) is a medium-chain saturated fatty acid that serves as a substrate for mitochondrial β-oxidation^33^. Its reduced concentration may reflect impaired fatty acid catabolism or peroxisomal compensation in CHB + MASLD. Like aspartic acid, caproic acid also showed a positive correlation with HDL-C, suggesting its potential involvement in maintaining hepatic lipid homeostasis. Beyond energy metabolism, medium-chain fatty acids such as caproic acid have been implicated in gut-liver axis regulation and anti-inflammatory responses^34^. Glycolaldehyde is a reactive aldehyde derived from glycolytic intermediates or lipid peroxidation^35^. Although typically associated with oxidative stress, its decreased levels in CHB + MASLD may reflect altered redox balance or downregulated glycolytic throughput under mitochondrial dysfunction. Clinically, glycolaldehyde was positively correlated with HDL-C and negatively with liver enzymes(ALT/AST) and BMI, suggesting it may serve as a marker of intact energy flux and metabolic resilience. Monomenthyl succinate is a methylated derivative of succinic acid, potentially formed during metabolic overflow or under hypoxic conditions. Its depletion may indicate dysregulated succinate metabolism, reflecting a shift toward reductive pathways or HIF-1α-mediated adaptation^36,37^. While less characterized in hepatic disease, its strong association with multiple clinical indices underscores its potential as a surrogate marker of mitochondrial metabolic efficiency. D-serine is a non-essential amino acid with neuromodulatory functions^38^. Elevated levels in CHB + MASLD may reflect dysregulated amino acid turnover, possible contributions from renal–hepatic interactions, or alterations in oxidative stress buffering. Succinic acid, a key intermediate in the TCA cycle, was significantly elevated in CHB + MASLD^39^. Beyond its metabolic role, succinate acts as a signaling molecule that activates HIF-1α, promoting inflammation and fibrosis under hypoxic conditions^27,40–42^. The enrichment of succinate in both the global and refined KEGG pathway analyses further highlights its central role in disease progression. Sebacic acid, a dicarboxylic acid derived from ω-oxidation of medium-chain fatty acids, was also consistently elevated in CHB + MASLD. Sebacic acid has been implicated in oxidative stress, inflammation, and fibrosis^43–45^, possibly via lipid peroxidation and dicarboxylic acid accumulation within hepatocytes. Its robust discriminatory power and pathophysiological relevance make it a strong candidate biomarker and a potential indicator of maladaptive lipid handling in the setting of metabolic dysfunction. While the causality of this association remains to be fully elucidated, the correlation suggests a close link between sebacic acid accumulation and hepatic metabolic stress.

In summary, these discriminatory metabolites converge on pathways governing mitochondrial energy production, lipid oxidation, and amino acid–redox balance, providing mechanistic insight into the metabolic complexity of CHB + MASLD. Their consistent importance across multiple algorithms, together with strong clinical correlations, supports their potential utility as both pathophysiological markers and components of predictive diagnostic models.

### Diagnostic Model Performance and Clinical Translation Potential

Building upon the machine learning–prioritized metabolite panel, we constructed a parsimonious yet highly discriminative diagnostic model integrating key serum metabolites and routine clinical indicators. The model demonstrated excellent performance in both the training and independent test cohorts, with AUCs exceeding 0.90, significantly outperforming individual clinical parameters. This suggests strong generalizability and supports its potential utility as a non-invasive screening tool for MASLD in patients with chronic hepatitis B (CHB).

Given that MASLD often coexists silently in CHB patients and may be underdiagnosed by conventional imaging, especially in early or non-obese cases, this metabolic model offers clinically actionable value^46^. Incorporating selected metabolites into CHB management protocols could improve risk stratification and early intervention, especially in resource-limited or imaging-inaccessible settings.

Beyond diagnostic performance, this study contributes novel insights and methodological strengths compared to previous MASLD metabolomics studies:Multi-algorithmic feature selection: By integrating three machine learning approaches (XGBoost, SVM, LASSO), we minimized model-specific bias and ensured robustness and reproducibility of the selected metabolic signatures.Clinical-metabolic correlation analysis: We systematically evaluated the associations between discriminatory metabolites and key clinical indicators (e.g., liver enzymes, lipid profiles), offering a foundation for developing combined metabolic-clinical risk scores.

Importantly, while MASLD-related metabolomics studies are emerging, research specifically targeting CHB-complicated MASLD remains scarce, particularly at the systems metabolic level. Our work addresses this gap and lays a conceptual and methodological foundation for future studies exploring metabolic mechanisms, disease progression, and treatment monitoring in this under-recognized subpopulation.

Looking forward, targeted validation of the identified metabolites via small-scale LC-MS/MS assays could facilitate the development of a clinical diagnostic kit. Such tools may ultimately enable routine MASLD screening in CHB patients, improving early detection, timely management, and long-term outcomes.

## Conclusion

In summary, our study reveals that CHB-related MASLD is underpinned by a distinct pattern of metabolic remodeling involving mitochondrial dysfunction, lipid oxidative imbalance, amino acid reprogramming, and disrupted redox homeostasis. Rather than isolated abnormalities, these alterations reflect an integrated metabolic response to chronic viral inflammation, hepatic lipid overload, and energy stress. The prioritized metabolites such as succinic acid, L-aspartic acid, caproic acid, and glycolaldehyde not only serve as effective diagnostic classifiers but also illuminate key metabolic inflection points in disease progression. This systems-level metabolic reprogramming offers a framework for future mechanistic studies and supports the translation of metabolite-informed models into non-invasive diagnostic tools for early MASLD detection in CHB patients.

## Acknowledgements

Not Applicable.

## Availability of data and materials

The data generated in the present study may be requested from the corresponding author.

## Authors’ contributions

Conceptualization, **Chuyang Wang**; Methodology, **Chuyang Wang**; Software, **Chuyang Wang**; Validation, **Chuyang Wang** and **Yutao Chen**; Formal analysis, **Chuyang Wang** and **Yutao Chen**; Investigation, **Chuyang Wang** and **Yutao Chen**; Resources, **Huanming Xiao**, **Jianxiong Cai**, **Ruihua Wang**, and **Wofeng Liu**; Data curation, **Xuan Zeng**; Writing—original draft preparation, **Chuyang Wang**; Writing— review and editing, **Qubo Chen** and **Xiaoling Chi**; Visualization, **Ming Lin**; Supervision, **Qubo Chen** and **Xiaoling Chi**; Project administration, **Qubo Chen** and **Xiaoling Chi**; Funding acquisition, **Qubo Chen**. All authors have read and agreed to the published version of the manuscript.

## Ethics approval and consent to participate

All participants provided written informed consent, and the study was approved by the Ethics Committee of Guangdong Provincial Hospital of Traditional Chinese Medicine, ZE2023-194-01.

## Patient consent for publication

Not applicable, as this study does not contain any individual person’s identifiable data.

**STable 1.**Contribution of metabolites to classification accuracy in machine learning.

